# Neuronal development shapes activity-dependent gene expression in a stimulus-specific manner

**DOI:** 10.1101/2025.02.19.638694

**Authors:** Jeronimo Lukin, Maximiliano S Beckel, Olivia Pedroncini, Sebastian A Giusti, Ivana Marcela Linenberg, Ines Lucia Patop, Ariel Chernomoretz, Antonia Marin-Burgin, Sebastian Kadener, Damian Refojo

## Abstract

Neuronal activity-dependent gene expression is fundamental to a wide variety of brain functions. However, how development progress and stimulation modalities specifically affect neuron transcription is not fully understood. In this work, we first investigate the influence of development on neuronal firing and activity-driven transcription. We used an RNA sequencing approach over 7 days *in vitro* (DIV) or mature 21 DIV neurons, comparing neuronal depolarization with potassium chloride (KCl) versus Biccuculine application, a synaptic modality to induce neuronal activity. To further investigate how different activity patterns influence gene transcription in mature neurons, we compared global gene expression in neurons treated with three different and extensively used activation protocols: KCl, Bicuculine (Bic), and TTX withdrawal (TTXw). Our results demonstrate a strong influence of development on activity-dependent gene expression, and showed that different patterns of neuronal activity induce different transcriptional profiles and exhibit distinct temporal dynamics for the same genes. These findings offer novel insights into the complex relationship between neuronal activity and gene expression, shedding light on the context-dependent nature of activity-dependent transcriptional responses.

## INTRODUCTION

Activity-dependent gene expression is a molecular mechanism in neurons crucial for multiple cognitive, sensory-motor, developmental, and emotional adaptive processes (Benito et al., 2011; Yap and Greenberg, 2018). Immediate early genes (IEGs) are rapidly induced upon neuronal stimulation in a protein synthesis-independent manner (Greenberg et al., 1985; Morgan et al., 1987; Fowler et al., 2011). Numerous studies have demonstrated that various external stimuli activate the expression of IEGs, including growth factors (Cochran et al., 1983; Cole et al., 1989), neurotrophins (Ghosh and Greenberg, 1995; Martinowich et al., 2003), glutamate (Bading et al., 1993), NMDA (Pappas et al., 2012), electrical stimulations (Lee et al., 2017), potassium chloride (KCl)-induced depolarization (Kim et al., 2010; Benito et al., 2011; Malik et al., 2014; Tyssowski et al., 2018; Carullo et al., 2020), Bicuculline (Bic) (Bas-Orth et al., 2017), and Tetrodotoxin withdrawal (TTXw) (Saha et al., 2011). From these studies, specific IEGs, such as Fos, Arc, Npas4, Homer1, Igf1, and Bdnf, have been further characterized for their roles in neuronal function and plasticity (Barth et al., 2004; Rial Verde et al., 2006; Hu et al., 2010; Shepherd and Bear, 2011; Bateup et al., 2013; Jakkamsetti et al., 2013; Spiegel et al., 2014; Joo et al., 2016; Mardinly et al., 2016; Sun and Lin, 2016; Diering et al., 2017). Importantly, not all IEGs respond to every activation stimulus, and the mechanisms underlying these differences are not yet fully understood.

The impact of electrical activity on neuronal development and connectivity is well documented, both in cell culture and *in vivo* paradigms (Shatz, 1990; Fields and Nelson, 1992; Goodman and Shatz, 1993; Spitzer, 2006; Valor et al., 2007; Andreae and Burrone, 2014; Martens et al., 2016). However, there is a significant knowledge gap in systematic research regarding how activity-dependent gene expression programs vary at different stages of maturation. Much of our knowledge about IEG mechanisms originates from studies conducted on primary neuron cultures stimulated at 7-10 days *in vitro* (DIV). However, at this developmental stage, neurons are not fully mature, raising the question about the influence of neuronal development on activation-induced gene expression.

The temporality of neuronal activation is another critical factor for shaping activity-dependent gene expression programs. While increased frequencies of electrical stimulation correlate with increased expression levels of c-fos (Sheng et al., 1993), the duration of neuronal activation also plays a significant role in shaping gene expression (Lee et al., 2017; Tyssowski et al., 2018). Moreover, specific bursting patterns of neuronal activation exert differential effects over gene expression profiles, even when neurons receive the same number of electric stimulations (Sheng et al., 1993; Lee et al., 2017; Iacobas et al., 2019). However, these studies utilized prolonged activation protocols (up to 5h), potentially confounding interpretation due to homeostatic phenomena. Additionally, these studies were performed in dorsal root ganglion (DRG) neurons, which remain silent in culture and lack dendrites and synaptic contacts. Consequently, the impact of acute activation patterns on gene expression in cultured cortical neurons needs to be further elucidated.

Here, we investigate the influence of neuronal development on the transcriptional response to activity by conducting a comparative analysis of the activity-induced gene expression between immature (7DIV) and fully differentiated (21DIV) neurons in culture stimulated by synaptic activity (Bic) and massive depolarization (KCl). Additionally, to address how different activity patterns influence gene transcription in neurons, we performed a comparative analysis of global gene expression in neurons acutely activated with KCl, Bic, and TTX withdrawal. Overall, our findings suggest that the transcriptional response to activity-driven stimulation is strongly influenced by the degree of neuronal maturation and that different modalities of neuronal activity elicit specific transcriptional programs with unique and distinct temporal dynamics.

## RESULTS

### The maturational stage of neurons influences neuronal firing and dictates the course of activity-driven transcription

Several studies have shown how basal transcriptional profiles change as neurons develop and their synaptic contacts increase and mature (Martens et al. 2016; Valor et al. 2007). However, how the degree of neuronal differentiation impacts the transcriptional programs induced by neuronal activity has not been systematically evaluated so far. To address this question, we performed an RNA-seq analysis after stimulating immature (7DIV) and mature neurons (21DIV) with two different activation modalities: synaptic activity by treating cells with Bic and membrane depolarization with KCl at different time points.

First, by evaluating 7DIV and 21DIV unstimulated neurons (Fig. 1A) we observed that neurons exhibit a more complex morphology and a higher synaptophysin expression, validating the expected increase of synaptic contacts as maturation progresses (Fig. 1B). Accordingly, whole-cell patch-clamp recordings indicate that neurons at 7DIV present a lower spontaneous firing rate that significantly increases at 21DIV (Fig. 1C), evidencing higher levels of network connectivity and basal activity in mature cultures. When performing a comparison of gene expression levels of stimulated and unstimulated neurons by RNA sequencing, the PCA analysis showed an apparent clustering of the samples belonging to different developmental stages (Fig. 1D). We observed 3.885 differentially expressed genes (DEG) between 21DIV and 7DIV neuronal basal gene expression (EV Table S1) with 2081 genes significantly up-regulated and 1804 genes down-regulated in 21DIV compared to 7DIV (Fig. 1E). Gene Ontology (GO) enrichment analysis on 21DIV vs 7DIV DEG revealed that the significantly enriched terms were related to the neuronal morphological maturation and synaptic organization and function (Fig. 1F). 572 of these 3885 DGE were mapped to unique SynGO annotated genes presenting a significant enrichment in synaptic cellular components (Fig. 1G).

**Figure 1:**
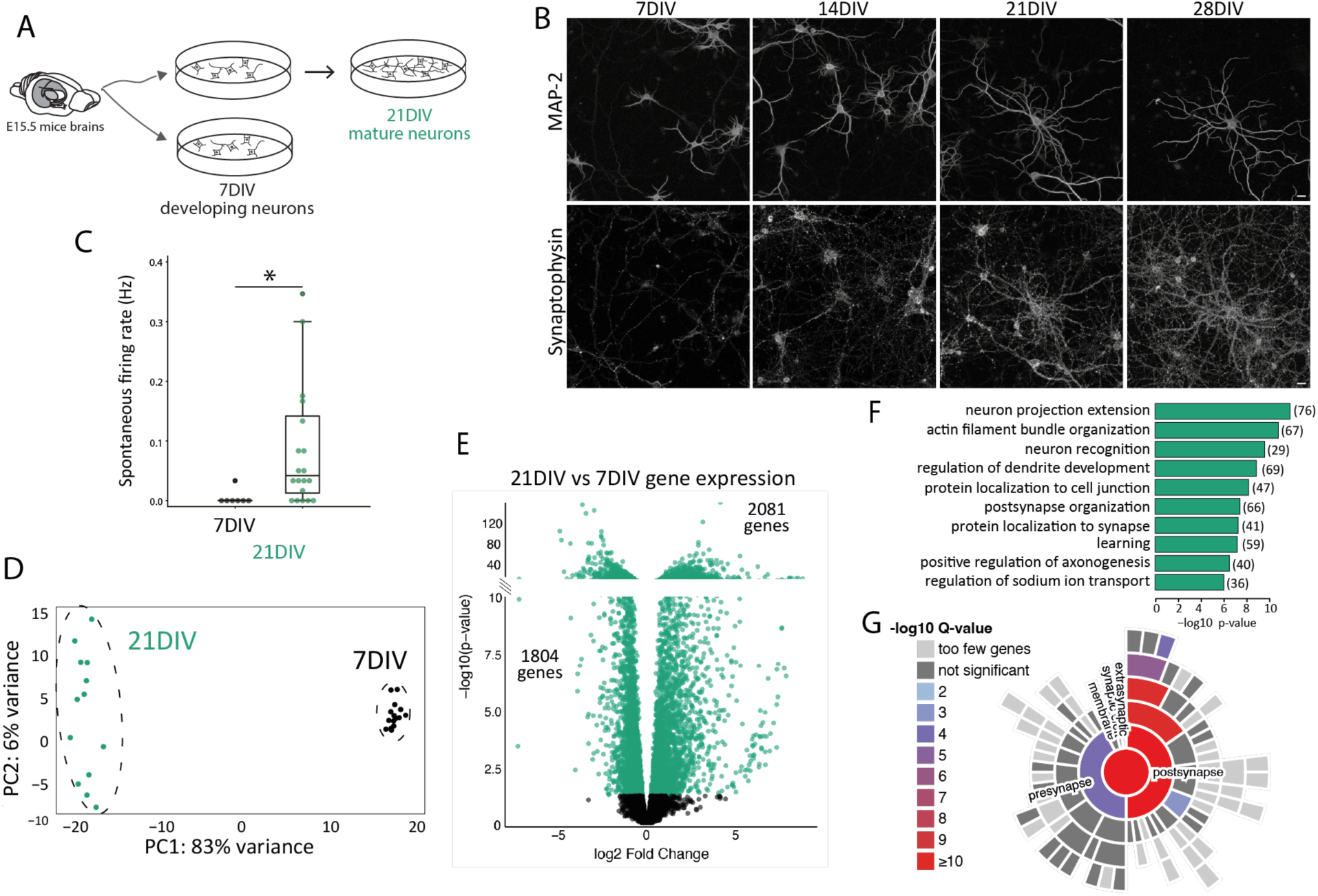
Developmental stage influences different neuronal molecular characteristics. A. Experimental design: primary cortical neurons from E15.5 mouse brains were cultured for 7DIV (developing) and 21DIV (mature). B. immunocytochemistry of MAP2 and Synaptophysin at different developmental stages. Scale bar = 20um C. Spontaneous firing rate measured by patch clamp recordings in 7DIV neurons (n=6) compared to 21DIV neurons (n=8). Each dot represents a single neuron measurement per well. Two-sample t-test: t-statistic = -2.548, p value =0.0194. D. Principal Component Analysis (PCA) of gene expression data from stimulated or unstimulated samples at 7DIV and 21DIV. E. Vulcano plot showing differentially Expressed Genes (DEG) 7DIV and 21DIV in basal conditions. The plot is divided, and the scale is adapted to accommodate the broad range of p-values. F. Gene ontology enrichment analysis. Enrichment in biological processes related to neuronal development and synapse function. The number of DEGs in each biological process is reported in brackets. G. SynGO enrichment plot: gene expression differences between 7DIV and 21DIV show an enrichment in genes related to synaptic cellular components.

To understand how these differences are translated into activity-driven transcription, we evaluated gene expression after 15min-and 180min stimulation with KCl and Bic followed by RNA sequencing (Fig. 2A). At 7DIV, KCl stimulation induced 433 DEG compared to unstimulated controls, while synaptic induction of neuronal activity using Bic only produced the induction of 39 DEG (EV Table S2). In mature neurons (21DIV), both types of stimulation induced high numbers of DEG: 1810 with KCl and 1245 after synaptic stimulation with Bic (EV Table S2). By comparing the DEG shared between stimulation conditions, we observed a large proportion of the genes exclusively induced in each condition, indicating specificity in the transcriptional response (Fig. C, EV Table S2). In addition, the proportion of shared DEG is higher between KCl/Bic in mature neurons than DEG shared upon each stimulation modality at different stages (Fig. 2C, EV Table S2). PCA also separated 7DIV and 21DIV samples depending on stimulation modality and time (Fig. 2D).

**Figure 2:**
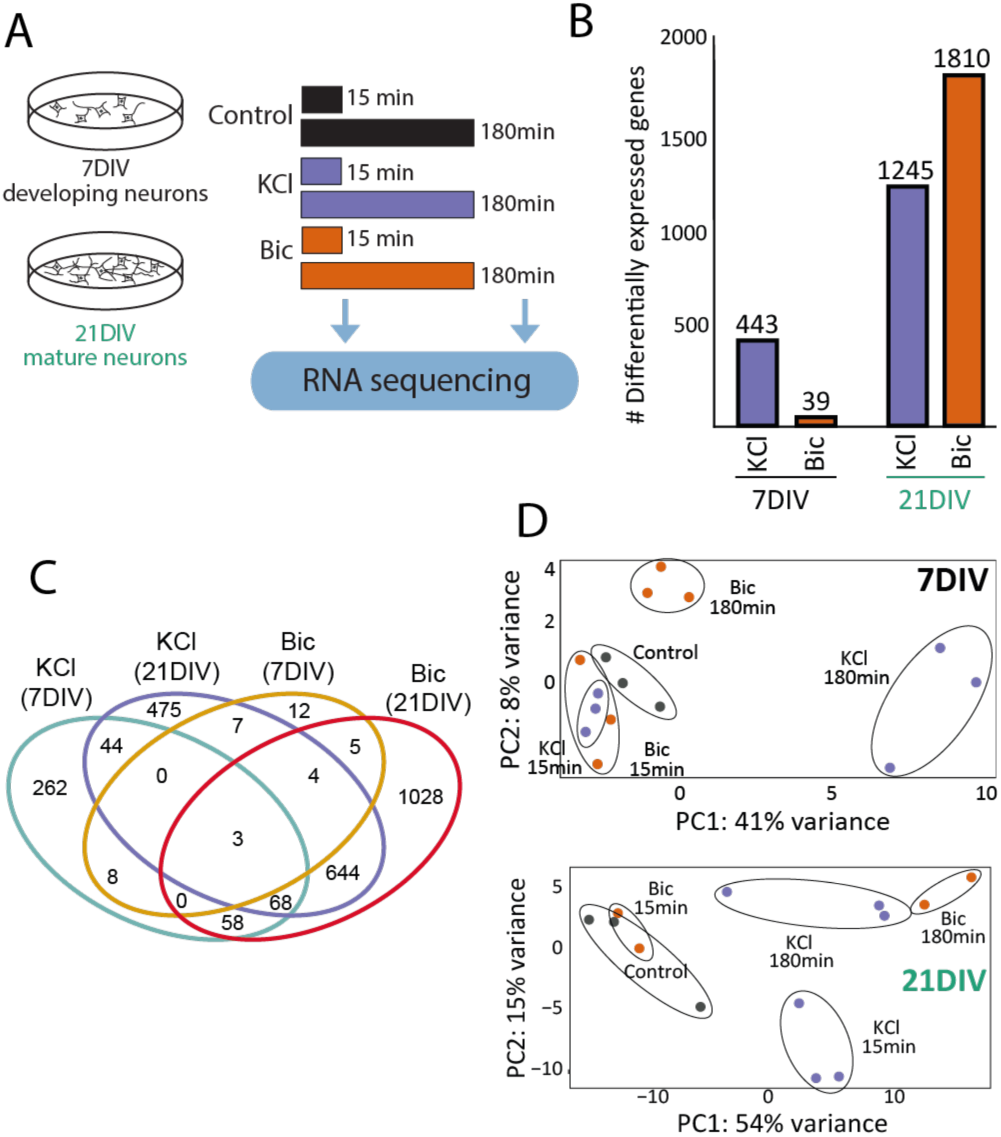
Activity-driven gene expression responses differences in 7DIV and 21DIV neurons. A. Experimental design: 21DIV neurons were stimulated with KCl or Bic for 15min or 180min, and RNA was extracted and sequenced to assess differential gene expression. B. Differentially expressed genes (DEG): number upon KCl of Bic stimulation at 7DIV and 21DIV. C. Venn diagram showing the number of overlapping DEG among experimental situations. D. PCA of gene expression data from 7DIV (top) or 21DIV (bottom) stimulated neurons shows separation based on stimulation time and modality.

The differences in activity-driven response might not be only due to the substantial molecular differences in the basal starting point (Fig. 1) but also due to the distinct ability of the neurons to physiologically respond to each stimulation at these two maturational time points. To investigate this, we performed patch-clamp recordings on cultured neurons and evaluated their electrical responses during the administration of KCl or Bic. At 7DIV, KCl application resulted in a massive and sustained depolarization (Fig. 3A). In contrast, Bic application did not alter neuronal activity, consistent with our previous observations of minimal changes in activity-dependent gene expression at this stage. The electrophysiological profile at 21DIV differed markedly from that at 7DIV. KCl application initially triggered a burst of action potentials (Fig. 3C), causing a transient increase in firing rate (Fig. 3F) and, in all cases, led to an irreversible shift in the membrane potential (Vm) towards a depolarized state. In contrast, Bic stimulation increased the firing rate in a subset of neurons (Fig. 3D, 3H) without inducing significant changes in Vm (Fig. 3G). These findings demonstrate that neuronal firing elicits distinct responses depending on the maturational stage. Altogether, these results highlight that neuronal maturation is associated with significant alterations in gene expression, particularly in genes involved in synaptic organization and function, which are likely linked to the observed changes in neuronal activity.

**Figure 3:**
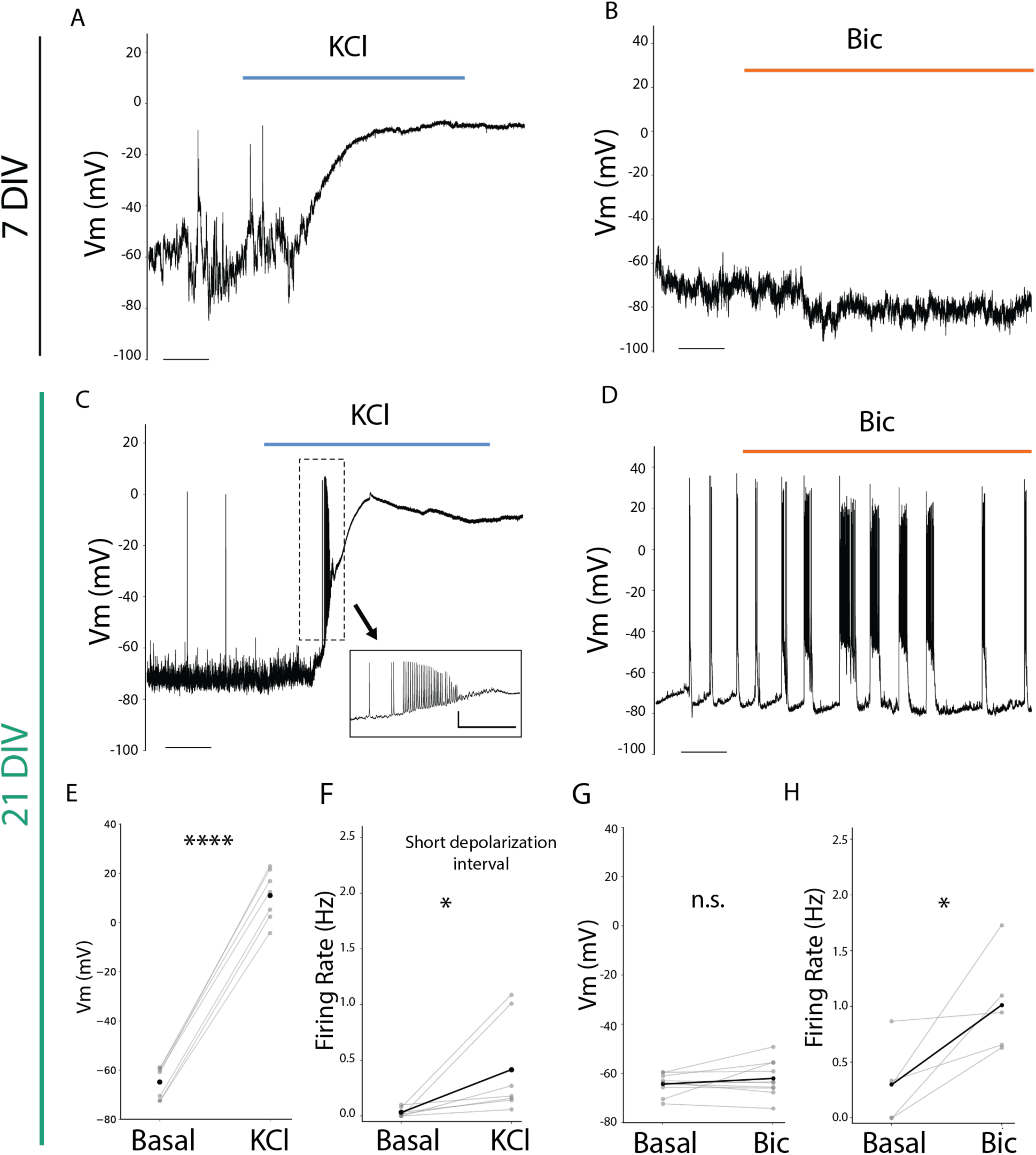
Electrophysiological response to KCl or Bic induction in 7DIV and 21DIV. A. Representative whole-cell patch-clamp recordings at 7DIV during KCl administration. Scale: t=50s. B. Representative whole-cell patch-clamp recordings at 7DIV during Bic administration. Scale: t=25s. C. Representative whole-cell patch-clamp recordings at 21DIV during KCl administration. Inset magnification illustrates a representative train of action potentials observed in these recordings at the beginning of depolarization. Scales: t=50s (main); and t= 5s; v=20mV (inset). D. Representative whole-cell patch-clamp recordings at 21DIV during BIC administration. Scale: t=25s. E. Voltage membrane (Vm) measurements of 21DIV neurons prior to and after KCl administration short window. t-statistic: -16.498, p= 1.30e-9; n=7. F. Firing rate (Hz) 21DIV neurons comparison between pre-stimuli and a short window during depolarization. t-statistic: -2.299498, p= 0.04e-9; n=7. G. Voltage membrane (Vm) measurements of 21DIV neurons prior to and after Bic administration. t-statistic: -1.207, p= 0.258; n=10. H. Firing rate (Hz) 21DIV neurons prior to and after Bic administration. t-statistic = -2.877, p= 0.045; n=5.

### Widely used stimulation protocols elicit specific and non-equivalent transcriptional responses

Building on the observed differences, we aimed to systematically analyze how the transcriptional profile and dynamics behave in mature neurons in response to acute stimulation by different stimuli. To investigate the temporal dynamics of gene expression, 21DIV neuronal cultures were exposed to KCl, Bic, or Tetrodotoxin removal (TTXw), and RNA sequencing was performed at three time points: 15, 45, and 180 minutes (Fig.. 4A). Although these methods are commonly used to activate neurons, their mechanisms of inducing activity actually differ significantly. PCA of these transcriptomic data showed a clear separation of samples exposed to TTXw from the KCl- and BIC-treated cells (Fig. 4B, upper panel). This result suggests that prior exposure to TTX for 48h induces a silencing activity that leads to a distinct transcriptional state compared to KCl or BIC, indicating that different activation modalities are not equivalent. When we performed a separate PCA for KCl and Bic (grouped) and TTXw, we observed differences in the temporal dynamics of transcription across the different stimulation conditions (Fig.. 4B, lower panels). A total of 2831 differential transcripts were found to be differentially expressed with respect to their basal condition: 1726 DEG upon TTXw, 1041 were induced by Bic, and 1036 after KCl membrane depolarization (Fig. 4C). Some DEG are shared among each group, with a greater proportion of genes shared between Bic and KCl when compared to TTXw (Fig. 4C). A heatmap of the 184 DEGs common to all three conditions further illustrates the distinct gene expression profiles and dynamics (Fig. 4D, EV Table S3). Collectively, these data showed that differences are related to which genes are differentially expressed in each situation and the temporal dynamics of gene expression induced by each neuronal activity modality.

**Figure 4:**
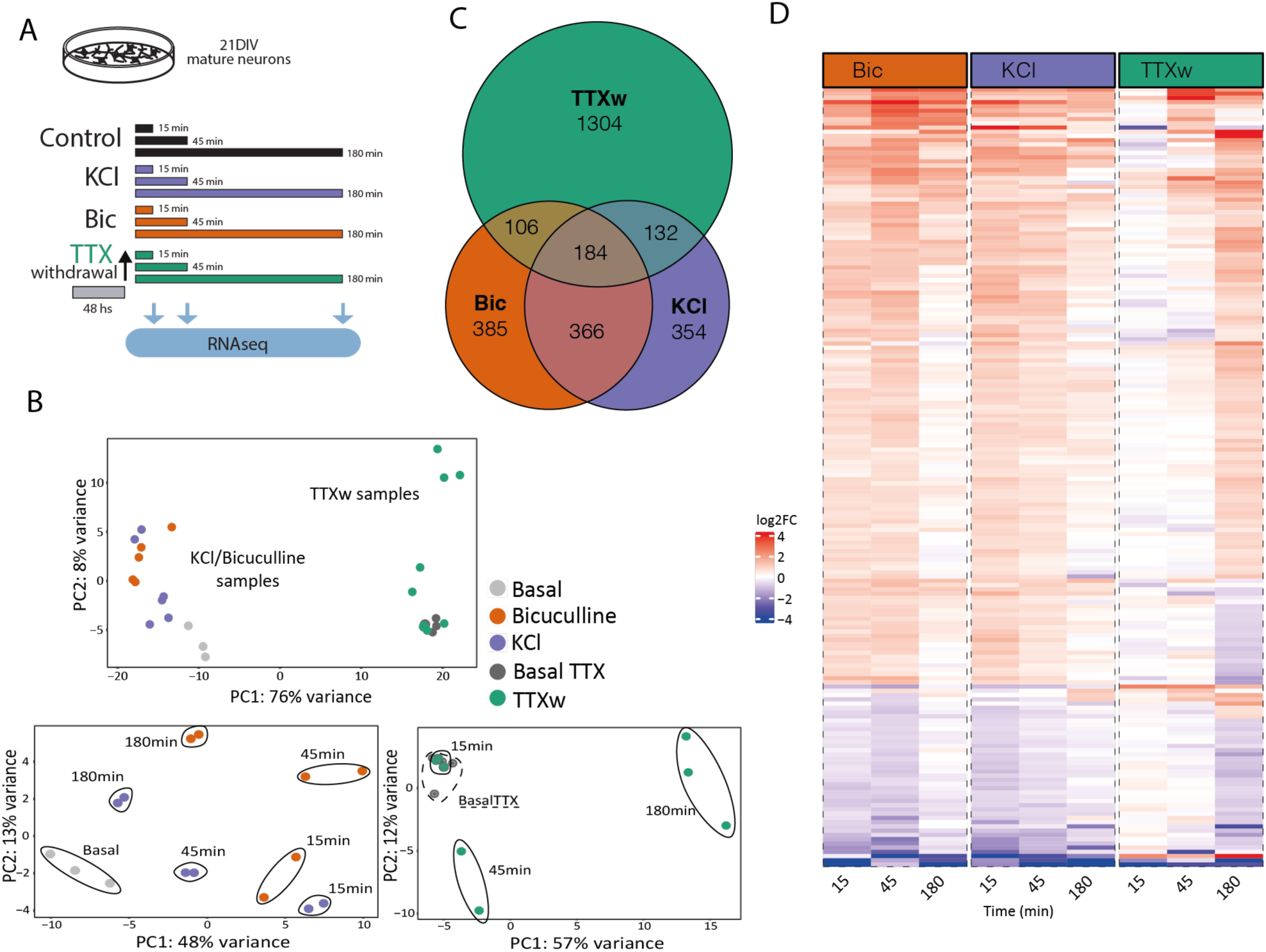
Gene expression programs induced by different neuronal activation modalities in mature neurons. A. Experimental design: Primary cortical neurons at 21DIV were stimulated with KCl, Bic, or TTXw for 15min, 45min, or 180min, and RNA was extracted and sequenced to assess differential gene expression. B. Gene expression PCA. Top: Samples (including non-stimulated controls) cluster into two groups: KCl and Bic vs. TTXw samples. Bottom: PCA of these two groups shows a separation based on stimulation duration. C. Venn diagram of DEGs among stimulation modalities. D. Heat map: fold induction levels (in log2) of the 184 shared DEG among stimulation modalities. Genes are grouped and ordered according to similar behaviors and dynamics.

To analyze immediate early genes’ expression dynamics, we selected 300 previously described IEGS by Kim et al. (2010) and Tyssowski et al. (2018) and performed a cross-comparison of the selected candidates among the different treatments. Out of these 300 IEGs, we observed a total of 161 DEG in response to some of the stimulation modalities: 35 of these genes were induced upon every stimulus, 74 genes were induced exclusively by one stimulus, and 54 genes had their expression levels altered by two of the protocols - 32 by Bic and KCl, 11 by Bic and TTxw, and 9 by KCl and TTXw- (Fig. 5A and 5B). 9 IEGs changed only upon KCl stimulation, 15 under BIC and 50 genes were exclusively modified by TTXw. The set of 35 shared IEGs whose expression levels changed in all treatments includes many of the most conspicuous and best-studied neuronal relevance IEGs such as the members of the Fos family (Fosb, Fos, Junb, Fosl2), Npas4, Arc, Bdnf, Nr4a2, Nr4a3, Egr3, Egr4, Gadd45b and Gadd45g, among others (Fig. 5C). Interestingly, we observed distinct expression dynamics for these shared genes depending on the stimulation modality (Fig. 5D). This collection of IEGs may represent a biologically relevant yet nonspecific gene transcriptional core that consistently responds to various activity-related stimuli, forming a basic ensemble of activity-responsive genes in neurons.

**Figure 5:**
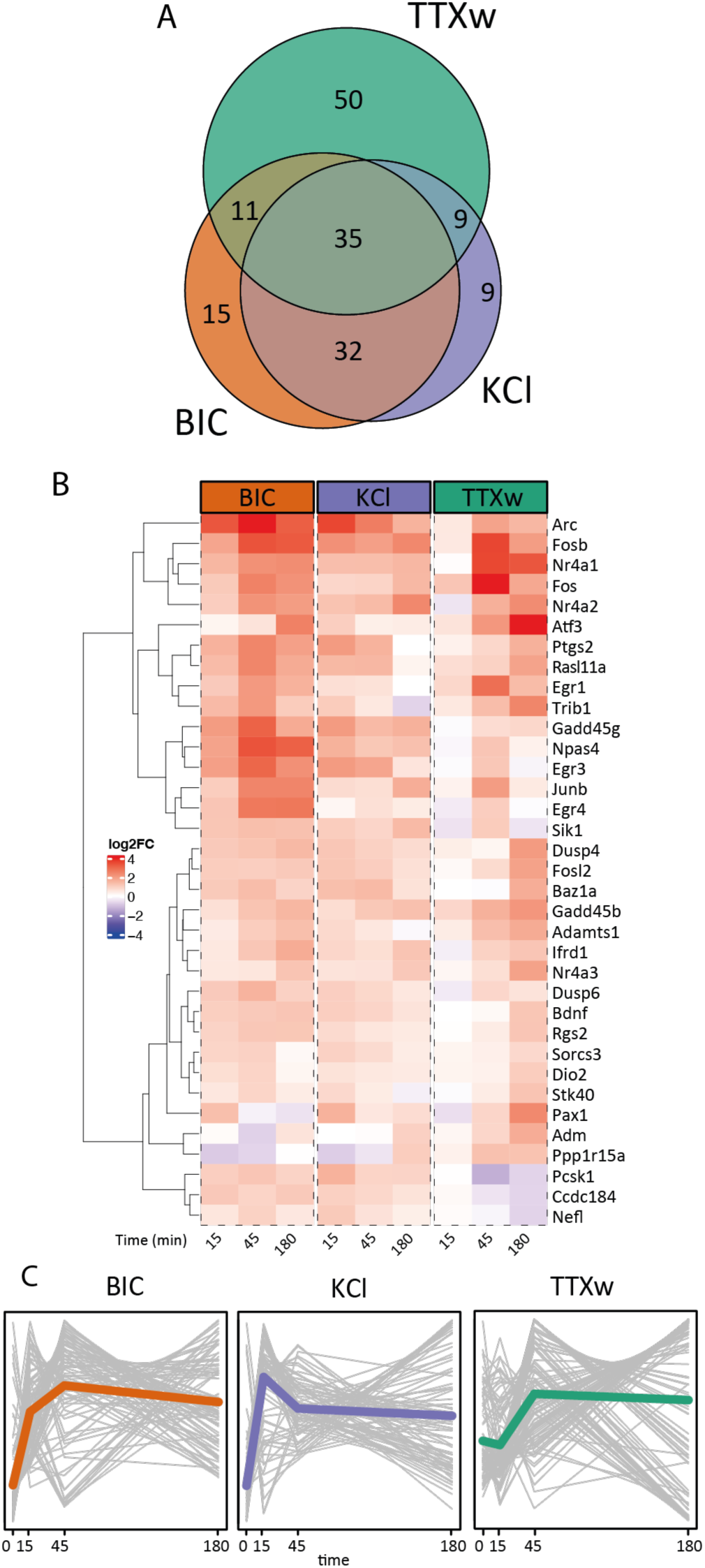
Immediate early gene expression dynamics. A. Venn diagram: overlapping IEGs differentially expressed among stimulation modalities. B. Heat map: fold induction levels (in log2) of 35 IEG differentially expressed upon stimulation modalities. Genes are grouped and ordered according to similar behaviors and dynamics. C. Average IEG expression dynamics. Each gray line represents a gene, and the colored line indicates the genes’ average trajectory corresponding to each dynamic.

### Analysis of gene expression dynamics analysis and GO characterization

We performed an additional analysis comparing DEG at each time point versus the basal gene expression values to better understand global gene expression dynamics among stimulation protocols. Notably, differences in the number and dynamics of DEG were observed among treatments: Bic generated a peak in the number of DEG after 45min; KCl depolarization induced a rapid gene induction of 925 DEG at 15min; and, upon TTXw, changes in the number of DEG were strongly observed only after 180min from drug withdrawal (Fig. 6A, EV Table S4). These results indicate that each stimulation protocol induces distinct temporal dynamics of gene expression.

**Figure 6:**
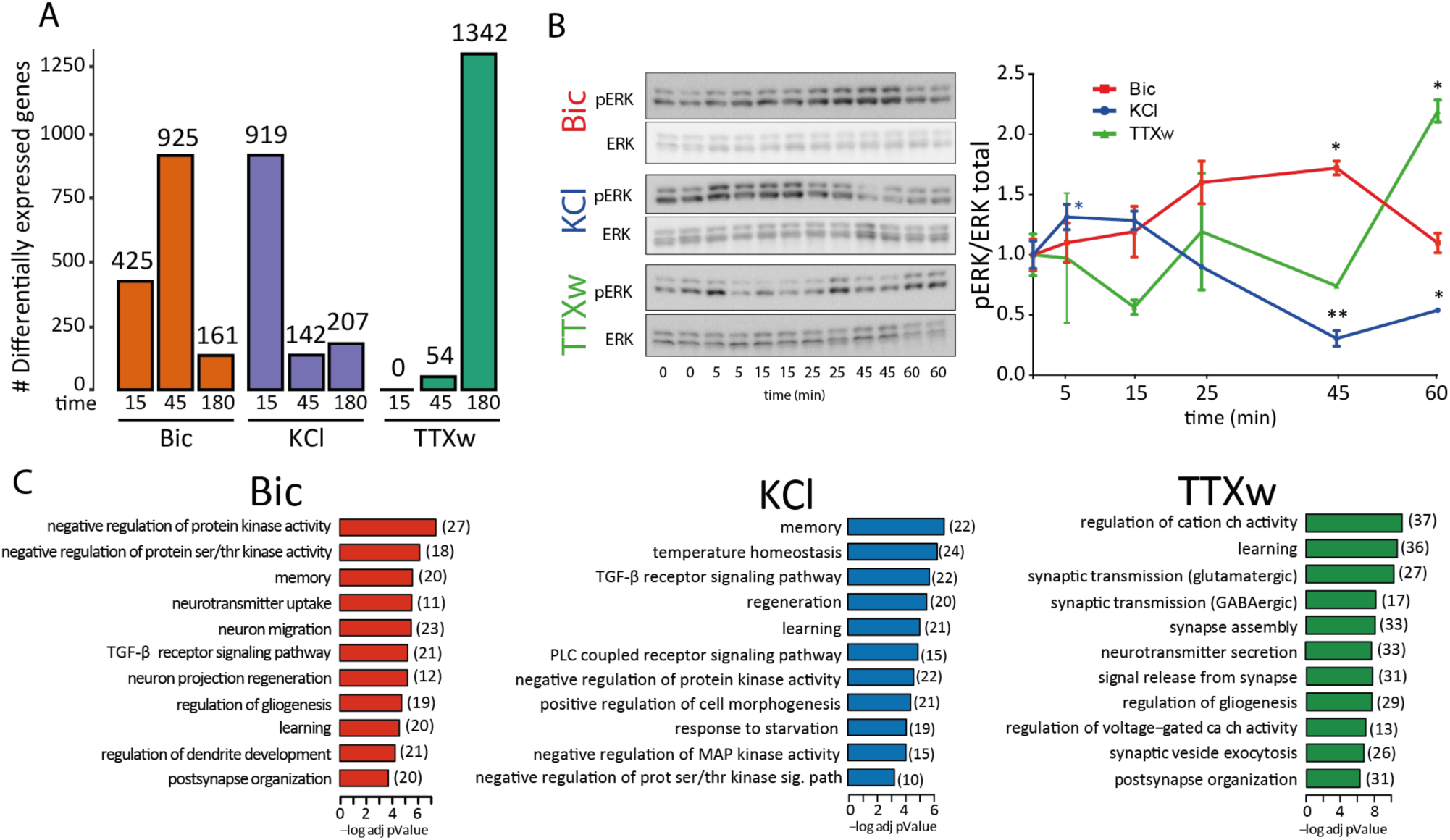
Gene expression dynamics analysis and ERK phosphorylation response to different stimuli. A. Bar graph showing the number of DEG at each time point. B. ERK phosphorylation kinetics upon stimulation treated with BIC, KCl, and TTXw for different times. Representative western blot for phosphorylated ERK (pERK) and total ERK. The dynamics of pERK relativized to total ERK are shown in the graph. Statistical comparison was made internally in each treatment, comparing with respect to the initial state at time zero. * p<0.05, ** p<0.01, one-way ANOVA followed by multiple comparisons via Dunnett’s test, n = 3. C. Gene ontology enrichment analysis. Biological processes enriched in Bic, KCl, and TTXw experimental groups are shown. In each stimulation modality, genes from the time point with more DEG were used for this comparison. The number of DEGs in each biological process is reported in brackets.

It has been previously shown that the ERK1/2-MAPK pathway is a key signaling hub orchestrating activity-induced gene-expression programs (Ha and Redmond, 2008; Tyssowski et al., 2018; Ohe et al., 2022). Thus, ERK1/2 phosphorylation can be used as a molecular proxi to assess the link between the induction methods and its downstream gene expression programs. Western blot analysis revealed ERK1/2 phosphorylation dynamics correspond with the peaks observed in gene expression upon each stimulation (Fig. 6B). Bic stimulation induced a significant increase in ERK1/2 phosphorylation at 45min (one-way ANOVA followed by multiple comparisons via Dunnet’s test: pERK1/2 Bic 45min vs. pERK1/2 Basal 0min, p=0.0455), KCl stimulation generated an initial increase in ERK1/2 phosphorylation at 5min, followed by a subsequent decrease (one-way ANOVA followed by multiple comparisons via Dunnet’s test: pERK1/2 KCl 5min vs. pERK1/2 Basal 0min, p=0.0494; pERK1/2 KCl 45min vs. pERK1/2 Basal 0min, p=0.0016; pERK1/2 KCl 60min vs. pERK1/2 Basal0min, p=0.0131), and TTXw resulted in a significant increase at 60min post-stimulation (one-way ANOVA followed by multiple comparisons via Dunnet’s test: pERK1/2 TTXw 60min vs. pERK1/2 Basal 0min, p=0.0482). Total ERK1/2 protein levels were largely unaffected. These findings indicate that each mode of neuronal activation triggers an ERK1/2 distinct activation pattern temporally aligned with the respective gene expression programs, suggesting that ERK1/2 phosphorylation might directly incluence the timing of downstream transcriptional responses.

Time-course analysis also allows us to differentiate between up-regulated and downregulated genes. Notably, the proportion of up-regulated to downregulated genes varies across time points (Fig. EV1A, EV Table S4). Interestingly, the time points with the highest number of DEG for each stimulus coincide with both the peak of up-regulated genes and the highest proportion of downregulated DEG. This result suggests a coordinated regulation of gene expression, where both upregulation and downregulation occur in a stimulus- and time-dependent manner.

To uncover the functional significance of the DEGs, we conducted a Gene Ontology (GO) functional enrichment analysis. Examining the enriched GO biological processes for each experimental condition, we found a direct relationship between the number of DEGs and the corresponding enriched GO terms (Fig. 6C, EV Table S5). Notably, following TTXw treatment, the enriched GO terms included a higher proportion of downregulated genes, as indicated by the z-score (Fig. EV1B). While general neuronal-related processes such as “learning” and “memory” were enriched across all treatments, specific neuronal and cellular GO terms were uniquely enriched depending on the stimulation modality (Fig. 6C, EV Table S5). Bic and TTXw treatments significantly enriched biological processes related to neuronal projections, dendritic projections, and synaptic organization. In contrast, these terms were not enriched following KCl treatment, which aligns with its mechanism of inducing a large influx of calcium without continuous synaptic activation. Overall, these findings further support the existence of stimulus-specific gene expression programs in neurons.

### Individual gene behavior and gene dynamic clustering

To gain deeper insight into individual gene temporal dynamics, we analyzed expression patterns at the single-gene level and identified three distinct groups with unique temporal profiles. The first group, including genes like Gadd45g, Arc, Ptgs2, and Rasl11a, exhibited expression patterns that aligned with the specific peaks for each stimulation modality (KCl at 15m, BIC at 45m, TTXw at 180m shown in Figure 6A & B) (Fig. 7A, first column). The second group, comprising genes such as Nr4a3, Egr4, Npas4, and AU023762, showed a similar response pattern to the three different activation treatments (Fig. 7A, second column). In contrast, genes of a third group, including Fos, Btg2, Junb, and Ccdc184, displayed distinct responses to each stimulus (Fig. 7A, third column). These results underscore the complexity and diversity of temporal gene expression dynamics in response to different neuronal activation modalities.

**Figure 7.**
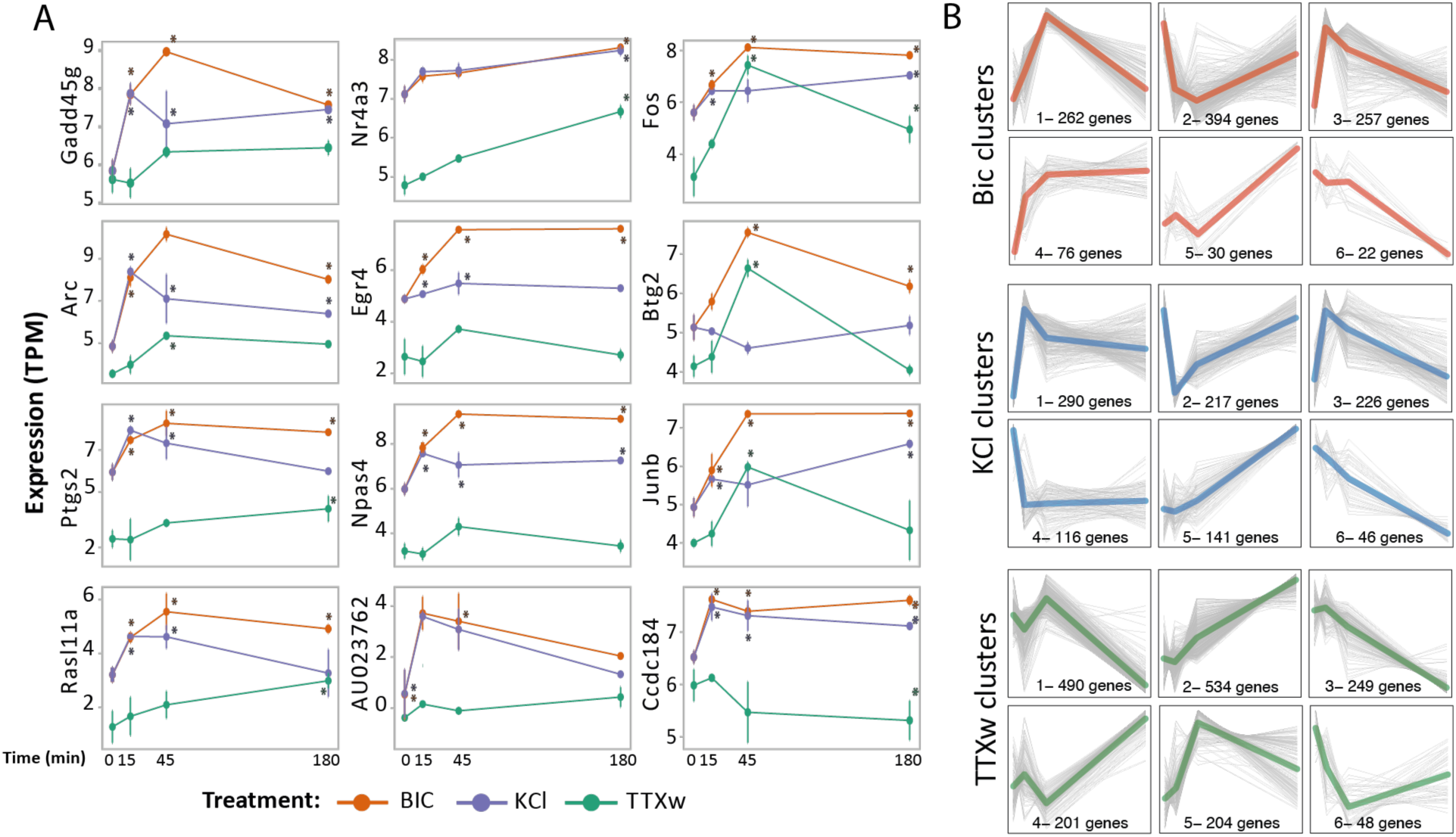
Individual gene behavior and gene dynamic clustering. A. Gene expression dynamics: selected genes’ expression levels are shown (TPM). Three gene groups with specific behaviors are presented: in the first column, transcripts peaking at the most prioritized time point for each stimulus (see Fig 5a); in the second column, genes responding similarly across the stimulation modalities.; and in the third column, genes behaving differently upon each stimulus. All presented genes exceed the statistical significance filters in their global dynamics. Asterisks indicate significance versus statistical measure of time-to-time comparison. B. Gene dynamics clustering of DEG upon each stimulation modality. The dynamic-tree cut method was used to infer clusters from the obtained hierarchical dendrogram and those clusters whose mean profiles had a distance less than 0.3 were merged. Each gray line represents a gene, and the colored line is the genes’ average trajectory corresponding to said dynamics. The number of genes that each of the 6 clusters of each treatment is indicated.

Using the dynamic tree-cut method, we performed gene clustering of DEGs based on their expression dynamic and identified six different gene expression dynamics associated with specific neuronal activity patterns (Fig. 7B, EV Table S 6). Most clusters exhibited a transient increase in gene expression with a peak at 15 or 45 minutes, followed by a return to initial levels (e.g., BIC_1 and BIC_3; KCl_1 and KCl_3; TTXw_1 and TTXw_5). In contrast, some clusters showed sustained changes in gene expression up to 180 minutes (e.g., BIC_4 and BIC_6, KCl_5 and KCl_6, TTXw_2 and TTXw_3). A third group of clusters displayed a marked change in gene expression direction at the intermediate time points (e.g., BIC_5, TTXw_1 and TTXw_4). Notably, clusters BIC_1 (peak at 45min), KCl_1 (peak at 15min), and TTX_2 (constant increase up to 180min) contained the highest proportion of genes among each treatment, consistent with the distribution of DEG observed at each time point and treatment (Fig. 6A). To evaluate gene responses and identify clusters of similar genes, we compare cluster composition along each stimulation condition (Fig. EV2). BIC_2/KCl_2 pair shares the most genes, with their dynamics closely aligned (Fig. 7B). Other pairs, such as BIC_3/KCl_1, BIC_3/KCl_3, BIC_1/KCl_1, and BIC_1/KCl_3, share 60–100 genes and display similar dynamics, highlighting substantial overlap between BIC and KCl treatments. TTXw clusters also overlap significantly with BIC and KCl, particularly TTXw_2, which shares genes with BIC_1, BIC_3, KCl_1, and KCl_3. However, TTXw_2 exhibits distinct dynamics, with sustained gene expression increases up to 180 minutes (Fig. EV 2A). Among the 184 DEGs shared across all activation paradigms, the most gene-rich combinations included BIC_1/KCl_1/TTXw_2, BIC_3/KCl_1/TTXw_2, BIC_1/KCl_3/TTXw_2, and BIC_3/KCl_3/TTXw_2 (Fig. EV2B). Genes with reduced expression were prominent in BIC_2/KCl_2/TTXw_3 and BIC_2/KCl_2/TTXw_1.Add (Fig. EV2B).

Our gene clustering analysis revealed distinct gene expression dynamics corresponding to specific neuronal activity patterns, providing valuable insights into the diversity of transcriptional responses to various modes of neuronal activation. These findings offer significant implications for understanding the regulatory mechanisms underlying neuronal stimuli responses.

## DISCUSSION

Previous research defined the timeline of synaptogenesis in primary neuronal cultures, showing that synapse formation begins around DIV7-12 (depending on culture conditions), reaching maturation 6-10 days later (Harris et al., 1992; Papa et al., 1995; Benson and Cohen, 1996; Li and Sheng, 2003; Kaech and Banker, 2006; Banker, 2018). In fact, not only the structure but also the functional matching between the pre- and postsynaptic compartments increases with neuronal maturation (Kay et al., 2011). The morphological and electrophysiological measurements of our cultured neurons indicate that they follow the same timeline (Fig. 1, 3). In particular, the absence of an evoked electrophysiological response to Bic-induced synaptic stimulation in DIV7 neurons (Fig. 3B), in contrast to the robust response in DIV21 neurons (Fig. 3D), suggests that functional synapses are largely absent at this developmental stage in our culture conditions. Accordingly, a comparative analysis of DIV7 vs DIV21 neurons revealed substantial transcriptional differences in basal gene expression, with 3.885 genes differentially expressed between the two populations (Fig. 1). Even though DIV7 neurons remain largely unresponsive to (Bic-induced) synaptic transmission, they do respond to KCl, although with a smaller gene expression response (443 DEG at DIV7 vs 1245 DEG at DIV21) (Fig. 2). Interestingly, these differences are not only quantitative but also qualitative, as only 115 common genes are differentially expressed at both developmental stages (Fig. 2). KCl treatment bypasses synaptic transmission, directly triggering action potentials in each neuron. Thus, the reduced sensitivity to KCl in DIV7 neurons indicates that signaling cascades and transcrptional machinery necessary to translate activity into gene expression programs are still immature and not fully functional at this early stage.

These findings hold relevance for a vast research landscape of studies addressing activity-related phenomena where experiments involving primary neurons treated with KCl at DIV7-12 are abundant. The utilization of KCl treatment on neuronal cultures has been widely employed to induce membrane depolarization with varying concentrations and durations of treatment. This approach has been successful as a hypotheses generator and in identifying key immediate early genes (IEGs) (Bading et al., 1993; Flavell and Greenberg, 2008; Lin et al., 2008; Kim et al., 2010; Yun et al., 2013; Malik et al., 2014; Spiegel et al., 2014; Ataman et al., 2016; Mardinly et al., 2016; Stroud et al., 2017; Sharma et al., 2019; Boulting et al., 2021). Additionally, variations in this method affect whether the neuronal depolarization is translated into neuronal activity as an action potential (Wheeler et al., 2008; Rienecker et al., 2022). Therefore, our findings emphasize the necessity of considering neuronal developmental stages when interpreting activity-dependent gene expression and suggest that it may be prudent to standardize the use of more mature neurons in future research.

As mentioned in the introduction, different studies have explored activity-dependent gene expression in a genome-wide manner either in cell cultures (Kim et al., 2010; Benito et al., 2011) and/or different models like *Drosophila melanogaster* (Chen et al., 2016) and mice. *In vivo* studies have significantly advanced our understanding of IEG induction, employing diverse stimuli such as intense light exposure after keeping the animals in dark housing (Hrvatin et al., 2018; Tyssowski et al., 2018; Xu et al., 2021), the induction of epileptic-type electrical activity (Lacar et al., 2016; Fernandez-Albert et al., 2019), or stimuli associated with the administration of abuse drugs (Muniz et al., 2017; Mukherjee et al., 2018). These studies described a broad range of transcriptional responses and IEG induction. A comparative analysis of that data reveals the presence of a core set of genes consistently emerging across various stimuli, including Egr1, Egr4, Egr2, Npas4, Nr4a1, Dusp1, Fos, Arc, Jun, Fosb, Btg2, Atf3, Junb, Gadd45g, Nr4a2, Ier5, and Irs2. Notably, we identified a core set of 35 IEGs responsive to all three activation stimuli tested, encompassing many of these responsibly conserved genes (Fig. 5). This suggests the existence of a core transcriptional program governing the genetic response to neuronal activation, irrespective of the specific intracellular mechanisms involved. However, the activation of this core gene set do has a stimulus-dependent temporal dynamics, revealing a complex interplay between shared and specific signaling pathways. This implies that the temporal patterns of gene induction serve as a molecular fingerprint, encoding information about the underlying mechanisms driving neuronal activation.

Furthermore, neuronal activity to gene expression mechanisms can be conceptualized as a translation of digital signals (e.g. pulses/ discrete patterns) into analog intracellular signals. This process has been previously described in non-neuronal cells such as NIH 3T3 and PC12 cell lines showing that pulsatile stimulus over ligand or light-induced receptor activation can not only initiate distinct signaling cascades but also generate different activation dynamics of the same kinase pathway, such as ERK which subsequently controls IEG expression in those cell systems (Toettcher et al., 2013; Wilson et al., 2017; Ravindran et al., 2020; Ravindran et al., 2022). In fact, ERK is known to act as a central regulator of IEG expression in neurons (Tyssowski et al., 2018). Remarkably, ERK activation dynamics closely paralleled the temporality of activity-dependent gene expression observed across KCl, Bic,and TTXw stimulation (Fig. 5B), indicating that transcriptional responses are, at least in part, governed by the precise timing of early intracellular signaling events.

For years, KCl, Bic, and TTXw have been used almost interchangeably to induce neuronal activity. Recently, emerging evidence has also shown the divergent impacts of these stimuli on specific cellular phenomena. For instance, Bic and TTXw lead to distinct effects on nucleus-synapse transport (Ch’ng et al., 2012), while Bic, KCl, and TTXw induce different Arc transcriptional bursts (Das et al., 2018). However, despite their widespread use, no comprehensive comparison of these activation modalities has been conducted to elucidate their specific roles in activation-regulated transcription. Our analysis revealed marked disparities in the gene expression programs triggered by the different stimuli. This underscores the importance of considering Bic, KCl, and TTXw as non-equivalent, unique activation modalities to avoid misinterpretations and enhance reproducibility across different fields in neuroscience, studying diverse aspects of neuronal activation.

A more precise approach to dissecting the effects of activity patterns might be the use of electrodes providing direct electrical stimulation over neuronal cultures. Previous studies in dorsal root ganglion (DRG) neurons have shown that diverse bursting electrical stimulation patterns can induce specific gene expression profiles (Sheng et al., 1993; Lee et al., 2017; Iacobas et al., 2019). However, the magnitude of expression changes observed is relatively modest, possibly due to the extended activation protocols used (up to 5 hours). It is important to note that these studies were conducted in DRG neurons that develop in absence of synaptic contacts and remain silent in culture.

The difficulties in evaluating activity-dependent gene expression changes in response to short-duration controlled stimuli explain the predominance in this field of research of protocols that stimulate neuronal activity for longer durations like KCl, Bic, and TTXw with less control over the firing frequencies of neurons. Attempts to perform controlled-frequency stimulations were also performed using optogenetics in neuronal cultures. For example, few studies have demonstrated precise temporal control of IEG induction, such as Fos, by light in neurons expressing ChR2 (Schoenenberger et al., 2009). Furthermore, recent studies have demonstrated that in vitro neurons can reliably follow optogenetic stimulation frequencies up to 10Hz for several minutes, but fidelity declines at higher frequencies (Yang et al., 2024). However, optogenetic stimulation using light also has limitations. Notably, blue light alone can induce activity-dependent gene expression in neuronal cultures, even in the absence of channelrhodopsins in neuronal cultures (Tyssowski and Gray, 2019). These effects appear to result from phototoxic interactions with molecules present in neuronal culture media (Duke et al., 2020). To overcome this limitation, future studies could utilize electrical stimulation at specific frequencies of single pulses or bursts over varying time points using multi-electrode array (MEA) plates, in combination with pharmacological blockers of excitatory and inhibitory neurotransmission (e.g. APV-NBQZ or kynurenic acid plus picrotoxin). Such an approach would allow for the isolation of cell-intrinsic activation mechanisms while minimizing the contribution of network-level activity.

The advent of single-cell RNA sequencing, especially with anticipated improvements in sequencing depth, will be instrumental both in cultures and in vivo setups. In neuronal cultures, it would help to understand the magnitude and variability in responses among individual neurons in culture. In vivo studies combining neuronal labeling techniques with single-cell RNA sequencing will be crucial to pinpoint gene expression uniquely in neurons that were activated or incorporated into a memory engram (FOS+ or ARC+) in response to more physiological changes, such as exposure to a novel environment (Lacar et al., 2016) or engagement in a fear conditioning paradigm (Cho et al., 2016).

We also observed a substantial number of genes that were downregulated following neuronal activation, a phenomenon that remains largely understudied. One possible explanation is that this downregulation facilitates the upregulation of other genes essential for activity-driven neuronal changes. Although no specific strong shared identity emerged among the downregulated genes, the negative chromatin regulator Hdac11 (Liu et al., 2020) showed decreased expression following KCl, Bicuculline, and TTXw. Similarly, genes linked to autophagy, proteasome and endoplasmic-reticulum-associated degradation such as Derl3, Prss36, Atg2b, Ulk1, Ulk2 and Stbd1, were also downregulated. As were genes involved with mRNA degradation pathways, including Ufl1 and Piwil2, the latter of which functions with piRNA to methylate and silence gene targets (Rajasethupathy et al., 2012). These findings suggest a potential reduction in mRNA and protein degradation processes in neurons during activation. While we did not identify upregulated genes associated with transcriptional repression machinery, uncovering the role of this widespread gene downregulation offers an opportunity to deepen our understanding of complex regulatory mechanisms triggered by neuronal activity.

In summary, our study reveals that neuronal development significantly impacts the transcriptional response to neuronal activity. Using KCl, Bic, and TTXw to induce distinct neuronal activation patterns, we found that each stimulus elicits unique gene expression profiles in mature neurons. These findings emphasize the crucial need to consider both neuronal developmental stage and activation modality when exploring activity-dependent gene regulation.

## METHODS

### Mouse primary neuronal cultures

Neurons were dissected from embryonic day 16 to 17.5 (E16 -E17.5) CD1 embryos of mixed sex. Culture preparation was performed as previously described (Giusti et al., 2024). Briefly, forebrain (cortex and hippocampal) from CD1 mouse embryos were dissected and a neuronal suspension was prepared through Trypsin digestion and mechanical disruption of the tissue. Neurons were plated in 24 multi-well plates at a density of 80cells/mm2 (150.000 cells per well) and maintained in Neurobasal-A media (ThermoFisher) with 2% B27 and 0.5 mMGlutaMAX-I (ThermoFisher) at 37 °C and 5% CO2. CD1 mice were provided by our Specific Pathogen Free Animal Facility. All procedures were done in accordance with local regulations and the NRC Guide for the Care and Use of Laboratory Animals, followed at IBioBA-CONICET and approved by the local Institutional Animal Care and Use Committee (Protocol number 2020-02-NE) and were following the general guidelines of the National Institute of Health (NIH, USA).

### Stimulation protocols

Neurons were used between 19–23 DIV or at 7DIV when indicated. Stimulations were performed in neuronal growing media, and drugs were added at the indicated final concentration. No previous silencing was performed in any of the stimulation protocols. Massive membrane depolarization was achieved by applying 55 mM extracellular potassium chloride (KCl). We triggered neuronal activity by treating neurons with 50uM Bicuculline (Sigma) to induce synaptic stimulation. As indicated in the 7DIV -21 DIV comparison, we also added 2.5mM M 4-Aminopyridine (Sigma) to the 50uM Bicuculline treatment. Synaptic rebound was induced by performing TTX withdrawal(Rutherford et al., 1997; Saha et al., 2011). Cultures were treated with 1 uM TTX (Tocris) for 48hs, and the TTX was then washed out through seven exchanges of 1 ml of medium with fresh control medium. Control neurons were washed identically and processed in parallel. Results are shown in comparison to TTX control situation (silenced neurons TTX, 48hs). Each stimulation was sustained until the indicated time.

### RNA sequencing and analyisis

RNA from primary neuron cultures RNA was extracted using the RNeasy mini kit (QIAGEN) with in-column DNase treatment (QIAGEN) according to the manufacturer’s instructions. 15-25ng of RNA were used as input for preparing 3’ RNA sequencing libraries following CelSeq2 protocol (Hashimshony et al., 2016), changing the UMI to six bases. Sequencing was performed on Illumina NextSeq 500 system. Raw reads were aligned to Mus musculus genome (version mm10) using STAR (Dobin et al., 2013). Reads were quantified using End Sequence Analysis Toolkit (Gohr and Irimia, 2019) for 3’ RNA libraries. Differential gene expression analysis was performed with DESeq2 R package. Differential gene expression analysis was done with edgeR (Robinson et al., 2009). We excluded from the analysis genes that do not reach 5 CPMs in at least 2 samples. Reads were normalized by the trimmed mean method (TTM) for each time point before differential expression analysis. Significant differential expression used a cutoff of FDR < 0.05 and fold-change of at least ±1.5. Genes were grouped and ordered according to similar behaviors and dynamics through the Ward.D2 algorithm of the R package called Hierarchical Clustering when presenting gene expression levels in heatmaps.

### Gene Ontology enrichment analysis and gene clustering

Enrichment analyses were performed using the EnrichmentBrowser package (Geistlinger 2016). For the GO analysis, categories with a number of genes between 20 and 200 were considered. Those categories with FDR-adjusted p-value < 0.05 were considered significant.

Clustering of gene time profiles was made from a distance matrix calculated as one minus the correlation of gene expression values. Only genes that were significant in at least one time point were considered. We considered the dynamic-tree cut method (Langfelder, Zhang, & Horvath, 2008) to infer clusters from the obtained hierarchical dendrogram. In addition, a merging step was made to join the clusters whose mean profiles had a distance less than 0.3.

### Electrophysiological recordings

For electrophysiological recordings neuronal cultures were performed as described above, but cells were seeded on coverslips. Each coverslip was transferred to a chamber containing Artificial Cerebrospinal Fluid solution (ACSF) (in mM): 125 NaCl, 2.5 KCl, 2.3 NaH2PO4, 25 NaHCO3, 2 CaCl2, 1.3 MgCl2, 1.3 Na^+^ ascorbate, 3.1 Na^+^ pyruvate, and 10 dextrose (315 mOsm). Continuous bubbling with 95% O2/5% CO2 was administrated to the bath. Recordings were made in neurons of 7DIV and 21DIV. Whole cell current clamp recordings were performed using microelectrodes (4-6 MΩ) with an internal solution of potassium gluconate (in mM): 120 potassium gluconate, 4 MgCl2, 10 HEPES buffer, 0.1 EGTA, 5NaCl, 20KCl, 4ATP-tris, 0.3 GTP-tris, and 10 phosphocreatine (pH = 7.3; 290 mOsm). Recordings were obtained using Multiclamp 700B amplifiers (Molecular Devices), digitized and acquired at 20kHz on a desktop computer using pClamp10 software (Molecular Devices). Spontaneous activity was recorded in current clamp mode before and during the infusion of the drugs to the bath. For KCl treatment, the resulting firing rate was calculated during the short period of depolarization compared to the first 30seg 30 s of the recording in basal conditions. For Bic treatment, 45% of evaluated neurons were responsive to the treatment. and In those neurons, we compared a 40seg 40 s window after the neurons are exposed to Bic with the first 30seg 30 s of the recordings in basal conditions. When a drug was infused into the bath, In every stimulated measuments, only one neuron per coverslip was recorded to ensure that the neurons had not been previously exposed to had not beed previously stimulatedstimulation. Membrane capacitance and input resistance were obtained from current traces evoked by a hyperpolarizing step of 10 mV. Series resistance was typically 10 to 20 MΩ, and neurons were discarded if it exceeded 40 MΩ.

### Western Blots

Cells and tissue were lysed in RIPA buffer containing protease inhibitors (Roche, Mannheim, Germany). Protein samples were separated by 8–10% SDS-PAGE (Laemmli, 1970) and transferred to 0.2-μm PVDF membranes (Millipore, Darmstadt, Germany). Chemiluminescence signal was acquired in a ChemiDoc station (BioRad, Munich, Germany) and analyzed using Image J. For ERK and phosphoERK detection the following primary antibodies were used at a 1:1000 dilution: rabbit Phospho-p44/42 MAPK -Erk1/2-(Cell Signaling Technology #9101) and rabbit p44/42 MAPK -Erk1/2-(Cell Signaling Technology #4695). Secondary antibody anti-rabbit IgG, HRP-linked Antibody (Cell Signaling Technology #7074) was used at a 1:4000 dilution.

### Immunofluorescence staining

Immunofluorescence stainings were performed as previously described (Refojo et al., 2011). In brief, neuronal cultures were fixed with pre-warmed 4% Paraformaldehyde containing 5% Sucrose for 20 min at room temperature and then washed with PBS. Samples were permeabilized and blocked in 5% BSA (Sigma Aldrich), incubated with primary antibodies (overnight at 4°C), followed by Alexa dye-conjugated secondary antibodies (Invitrogen). Samples were mounted in VectaShield medium (Vector Laboratories). Primary antibodies used were used at a 1:100 dilituon: anti-Synaptophysin rabbit policlonal (Abcam, ab 14692) and anti MAP2 chicken (Abcam, ab 5392). Secondary antibodies were used 1:2000: Goat anti-rabbit IgG (H+L) Alexa Fluor 594 (Invitrogen, A-11037) and Goat anti-Chicken IgY (H+L), Alexa Fluor® 647 (Invitrogen, A-21449).

## Supporting information

EVTable1

EVTable2

EVTable3

EVTable4

EVTable5

EVTable6

## DATA AVAILABILITY

The bulk RNA-seq data generated in this study are publicly available at Gene Expression Omnibus (GEO) with accession number **GSE277512.** https://www.ncbi.nlm.nih.gov/geo/query/acc.cgi?acc=GSE277512

## ACKNOWLEDGMENTS

This work was supported by grants from Agencia Nacional de Promoción Científica y Tecnológica (grant 2017-4346 To SAG and 2019-03343 to DR), Consejo Nacional de Investigaciones Científicas y Técnicas (Fellowships and positions to JL, MB, OP, IML, SAG, AC, AMB, DR and grant PIP-11220210100755CO to SAG), Fondo para la Convergencia Estructural del Mercosur-FOCEM-grant COF 03/11 (DR), Max Planck Society grant (DR), Volkswagen Foundation (DR) and the Swiss National Science Foundation-SNSF, (DR and AMB).

Jeronimo Lukin current affiliation is Icahn School of Medicine at Mount Sinai, New York, USA. Olivia Pedroncini current affiliation is The Francis Crick Institute, London, United Kingdom. Ines Patop current affiliation is Harvard Medical School, Massachussets, USA.

## AUTHOR CONTRIBUTIONS

DR, AMB, SK, JL and SAG conceptualized the study and designed the experiments. JL, SAG, OP and IL conducted the experiments and collected the data. JL, MB, ILP, AC and SK performed the RNA sequencing conceptualization and analysis. JL, DR and AMB wrote the original draft, and all the other authors edited and approved the manuscript.

## EXTENDED VIEW FIGURE LEGENDS

**Extended View Figure 1:**
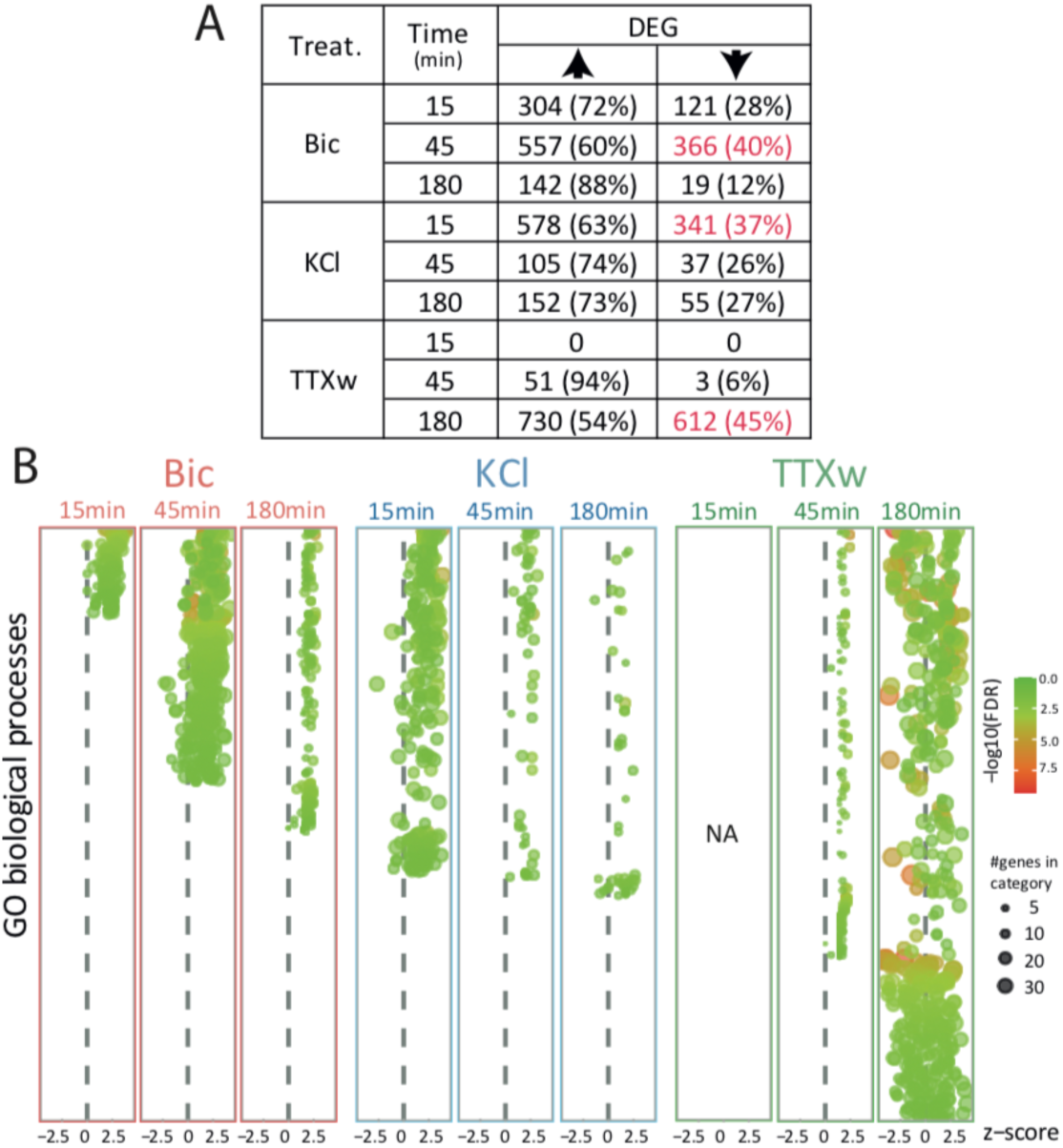
Gene expression dynamics analysis responses to different stimuli. A. Table presenting the number and percentage of DEG with significant increase or decrease in expression levels. B. Diagram showing the characteristics of GO biological processes found enriched in each experimental condition. Each circle represents a significantly enriched GO group. The color of the circles indicates the statistic value of that functional enrichment, while the size of the circles represents how many genes belonging to that group were found. Z-score estimates whether DEG present in each group are positively (z score > 0) or negatively (z score < 0) regulated compared to the basal situation.

**Extended View Figure 2:**
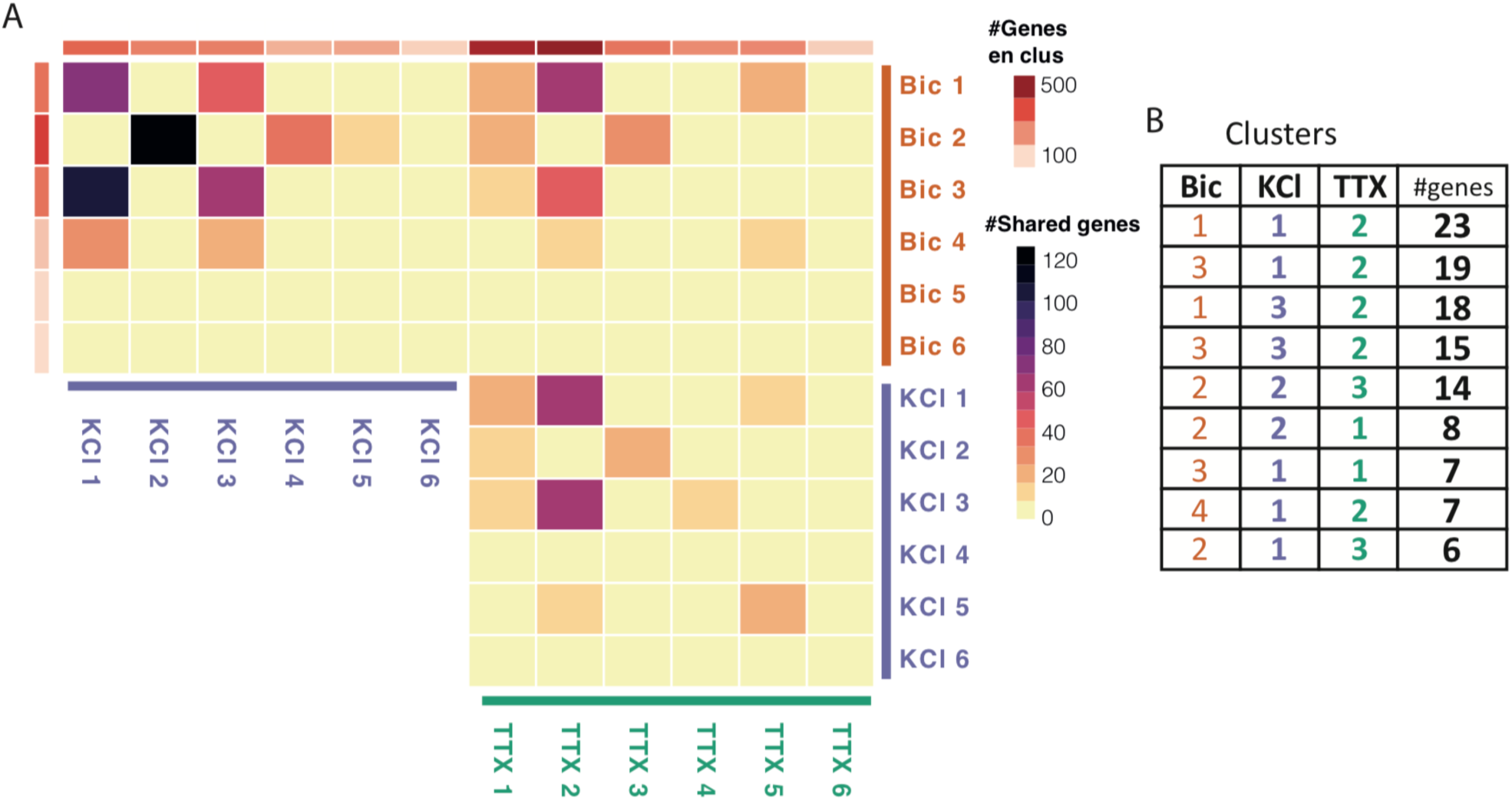
Gene clustering analysis. A. Distributions of the number of genes corresponding to each dynamic clusters and proportion of shared genes among clusters of different treatments. Darker squares correspond to pair of clusters that have a greater number of genes in common. The color of each row or column indicates the number of genes in that cluster at one end and the treatment to which they correspond at the other. B. Cluster trios with top amount of the 184 shared DEG upon Bic, KCl and TTXw.

